# Joint analysis of expression and variation at single cell resolution by scVar

**DOI:** 10.1101/2025.10.12.681753

**Authors:** Wei Zhao, Fei Yang, Wenting Zong, Xinchang Zheng, Yiming Bao

**Author notes:** **Corresponding author** Correspondence to Xinchang Zheng or Yiming Bao.

## Abstract

Tumor heterogeneity represents a critical determinant affecting both cancer diagnosis and therapeutic efficacy. Traditional bulk sequencing approaches are limited in their ability to resolve genomic alterations at the resolution of individual cells. In this study, we present scVar, an integrated analytical framework designed for mutation profiling using single-cell transcriptomic data. scVar enables sensitive and robust detection of single-nucleotide variants, particularly facilitating the identification of low-frequency variants. The framework incorporates customizable filtering parameters including variant allele frequency, and provides comprehensive functional and clinical annotations for identified mutations. Furthermore, scVar includes downstream analytical modules that facilitate the joint investigation of transcriptomic and mutational profiles within single cells. Benchmarking on simulated datasets and matched tumor samples demonstrates that scVar consistently outperforms existing methods, especially in detecting variants of low abundance, and shows strong concordance with whole-exome sequencing data. Application of scVar to non-small cell lung cancer samples effectively characterized ITH across spatially distinct tumor regions and diverse cellular populations. Collectively, scVar offers an integrated platform for concurrent analysis of somatic mutations and transcriptomic data derived from single-cell RNA sequencing, providing a powerful tool for elucidating tumor evolution and the complex interplay between genomic heterogeneity and the tumor microenvironment.

## Background

Cancer continues to represent a significant global health challenge, accounting for approximately 20 million new cases and 10 million deaths in 2022, as reported by the GLOBOCAN 2022 database (https://gco.iarc.fr/en). A major obstacle in the advancement of cancer therapeutics is the complexity of tumor heterogeneity, a multifactorial phenomenon that compromises diagnostic accuracy, therapeutic efficacy, and prognostic predictions[1]. Tumor heterogeneity is characterized by inter-tumor heterogeneity, which denotes the differences among tumors of the same type across different patients, and intra-tumor heterogeneity, which refers to the coexistence of multiple cancer cell subpopulations with distinct genotypes, phenotypes, and functions within a single tumor of the same patient. Intertumoral heterogeneity (ITH), compromises precise cancer diagnosis and effective therapy[2,3]. ITH has been identified as a prognostic factor for reduced survival across multiple cancer types[4] and a key driver of tumor evolution and therapeutic resistance[5–7]. Advances in high-throughput sequencing platforms, including whole-exome sequencing (WES) and whole-genome sequencing (WGS), have enabled quantitative characterization of tumor heterogeneity. These approaches facilitate the reconstruction of tumor clonal architectures and provide insights into evolutionary trajectories and mechanisms underlying drug resistance[8–10]. However, traditional bulk sequencing approaches lack single-cell resolution, limiting the accurate depiction of subclonal structures and constraining our understanding of the spatial and temporal dynamics of tumor heterogeneity.

The development of single-cell sequencing has made it possible to extend the study of genes and their functional alterations to the single-cell level[11]. Single-cell genome sequencing is the best way to identify somatic mutations in single cells. However, since each cell contains only two copies of DNA, single-cell DNA sequencing (scDNA-seq) faces many challenges, such as amplification bias and low coverage^9^. These can reduce the quality of scDNA-seq and increase the difficulty of mutation identification. Alternatively, somatic mutations can be detected directly from high-throughput single-cell assays, such as single-cell RNA sequencing (scRNA-seq). Notably, 10X Genomics scRNA-seq is widely adopted in transcriptome analyses due to its high throughput, cost efficiency, high mRNA detection sensitivity, and robust capacity to capture rare cell populations[12–14]. Furthermore, this technology has advanced our understanding of tumor biology by elucidating mechanisms underlying tumor heterogeneity, including clonal evolution dynamics[15,16], interactions within the immune microenvironment[17,18], and transcriptional plasticity among cellular subpopulations[19]. Importantly, scRNA-seq datasets contain latent genomic information, including somatic mutations, which remain underexploited due to methodological constraints[20].

Existing computational tools for mutation detection from 10X Genomics scRNA-seq data, including cellSNP[21], Monopogen[22], and SComatic[23], exhibit certain limitations. These include restricted accuracy stemming from the resolution of cell type annotations, insufficient sensitivity for detecting low-frequency mutations, and the lack of comprehensive software platforms that integrate single-cell transcriptomic data with mutation information.

To overcome these challenges, we developed scVar, a framework that extends mutation detection capabilities from bulk transcriptomic data to the single-cell level. scVar facilitates *de novo* identification of single-nucleotide variants (SNVs) from 10X Genomics scRNA-seq data without the necessity for paired normal samples or predefined cell-type annotations. The framework integrates seamlessly into standard 10X Genomics workflows and supports comprehensive single-cell transcriptome analysis, SNV detection, as well as downstream comprehensive analysis and visualization. This enables users to correlate transcriptional heterogeneity with genetic variation at single-cell resolution and to investigate the mutation landscape in details.

## Results

### Overview of scVar

scVar facilitates the comprehensive identification and analysis of somatic single nucleotide mutations from scRNA-seq data through a multi-layered analytical framework (Fig. 1). The workflow initiates with preprocessing and fundamental analysis of 10X scRNA-seq data, encompassing alignment, noise reduction, clustering, and cell-type annotation. The core stage of scVar involves mutation detection and annotation. During this phase, high-quality binary alignment map (BAM) files are reconstructed using cell barcodes, and candidate mutations are identified through GATK[24–26], optimized with parameters specifically adapted for mutation detection in 10X scRNA-seq datasets (Supplementary Note and Supplementary Fig. 1). Subsequently, genotype assignment is conducted at the single-cell level through parallelized computation, accompanied by noise reduction employing stringent filtering criteria. To reduce false positives, scVar applies rigorous filters, including the requirement that mutations be observed in at least two cells and the exclusion of common single nucleotide polymorphisms (SNPs) based on population frequency databases. Functional annotation of identified mutations is performed using the Ensembl Variant Effect Predictor (VEP)[27] and Annovar[28], supplemented by clinical relevance assessments referencing Cancer Hotspots[29,30], COSMIC[31], ClinVar[32], and other databases. Following this, scVar integrates mutation data with single-cell transcriptomic gene expression profiles into a MuData format file, thereby enabling streamlined and integrated downstream analyses. The analytical pipeline of scVar comprises three principal modules. The first module, mutation profiling and heterogeneity assessment, involves characterization of global and cell population-specific mutation signatures, evaluation of tumor mutational burden (TMB), assessment of clonal diversity using entropy and Simpson’s diversity index, and clonal clustering based on binary sparse mutation matrix. The second module focuses on functional annotation, encompassing the identification of cell population-specific mutations, enrichment analyses of oncogenic pathways and Gene Ontology (GO) terms, as well as pseudotime analysis to characterize the developmental states of mutated cells within the cellular differentiation trajectories. The third module provides statistical summaries such as estimation of cell type proportions, evaluation of alignment quality metrics, and visualization of the most recurrent mutations by mapping mutated cells onto Uniform Manifold Approximation and Projection (UMAP) embeddings with enhanced annotation.

**Figure 1.**
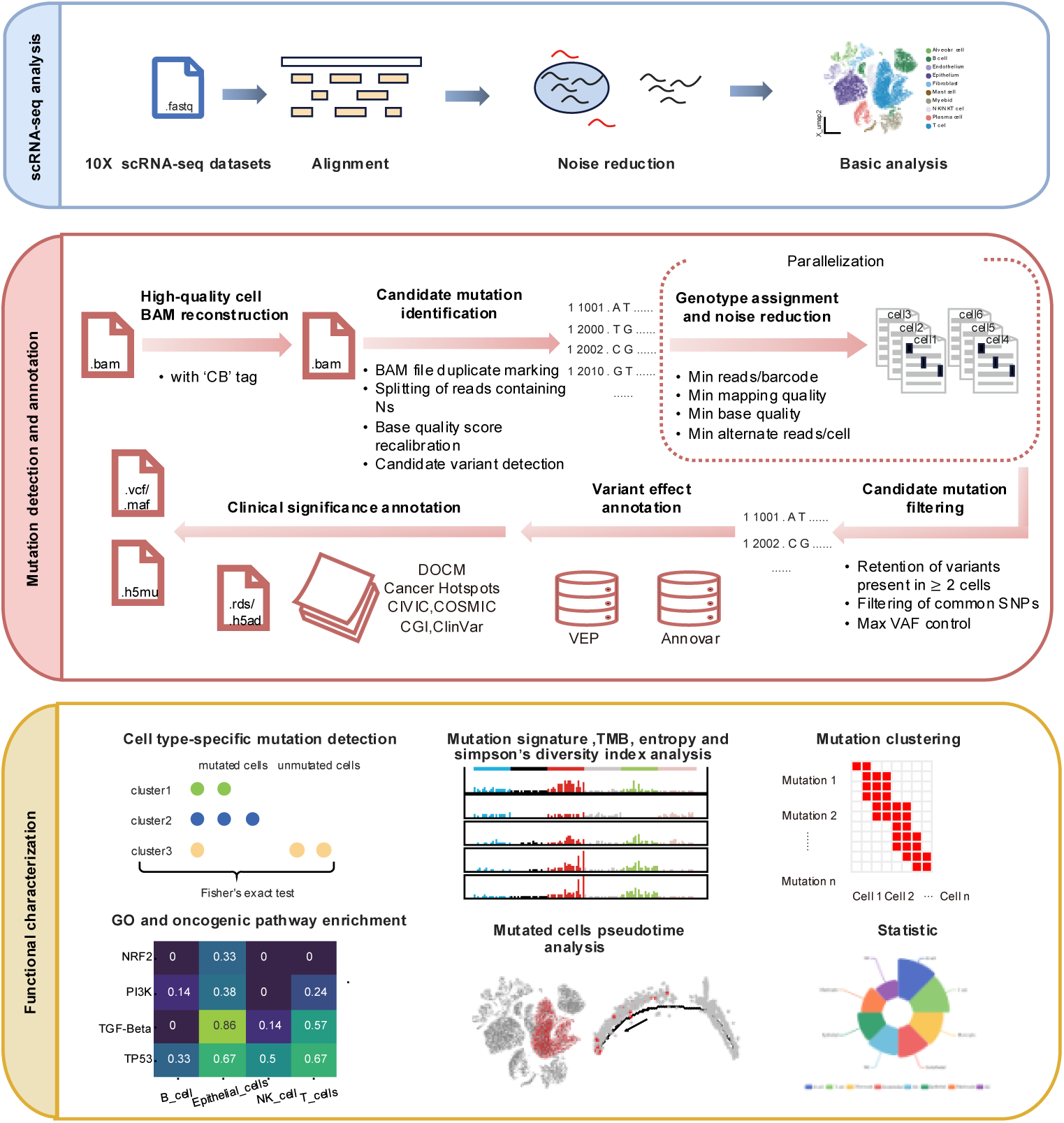
Overview of scVar. Schematic overview of the analytical workflow, illustrating the initial stages of data preprocessing, sequence alignment, quality control, fundamental single-cell transcriptomic analyses, and cell type annotation, followed by the principal steps of mutation processing, including somatic variant identification, genotyping, variant filtering, and functional annotation, and culminating in the integrated analytical modules implemented within the scVar framework.

In summary, scVar presents a comprehensive and integrated framework for the detection and analysis of SNVs derived from scRNA-seq data. It facilitates robust downstream analyses aimed at elucidating genomic heterogeneity, mutation dynamics, and cell-type-specific functional drivers. Furthermore, scVar can be seamlessly integrated into the standard 10X Genomics pipeline, thereby furnishing additional insights at the somatic mutation level.

### Performance evaluation of somatic mutation detection using simulated data

To establish a robust evaluation framework for assessing somatic mutation detection performance in 10X Genomics scRNA-seq data, we designed and implemented a comprehensive *in silico* spike-in experiment using five matched lung tissue samples.

These baseline datasets were preprocessed by globally reverting all detectable variants to reference alleles. High-confidence candidate loci were selected with at least 10X coverage. This was followed by *in silico* simulated mutations at controlled variant allele frequency (VAF) ranging from 10% to 50%, achieved by read-level modifications. Quantitative benchmarking against alternative single-cell variant callers—including cellSNP, Monopogen, and SComatic—revealed that scVar consistently outperformed these methods across all evaluated VAF thresholds. Specifically, scVar achieved the highest recall rates and demonstrated robust F1-scores, surpassing the performance of the second-best method, Monopogen. This performance advantage was especially notable at lower VAF thresholds, where scVar exhibited enhanced sensitivity in detecting rare somatic variants, thereby confirming its efficacy in resolving rare mutations from sparse transcriptomic data (Fig. 2a).

**Figure 2.**
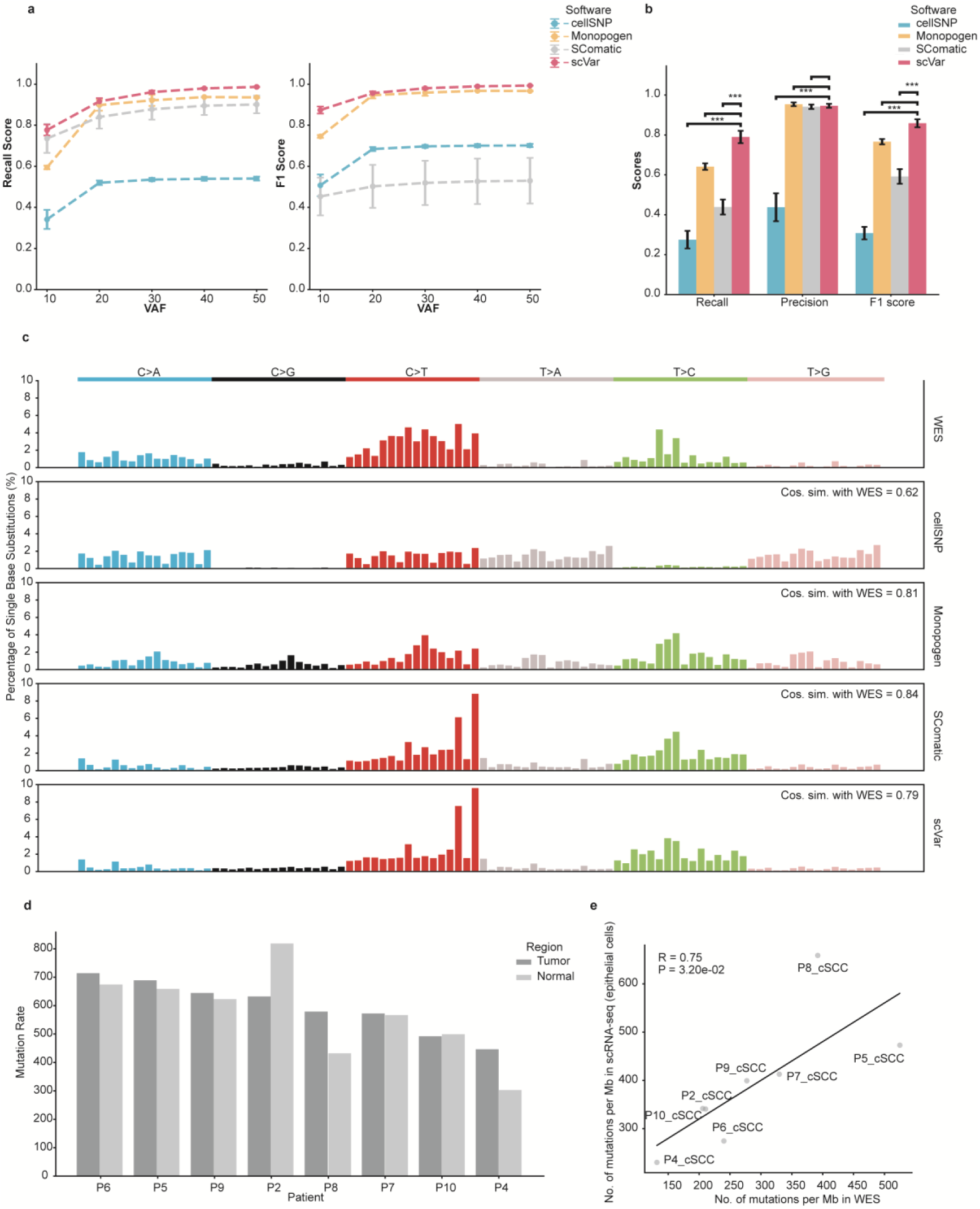
Comparison of the performance of scVar against other mutation detection methods. **a.** Performance of cellSNP, Monopogen, SComatic, scVar for the detection of somatic mutations in the simulation data. **b.** Performance of cellSNP, Monovar, SComatic, scVar for the detection of somatic mutations in the scRNA-seq data from cSCC. Significance with respect to scVar using the Student’s t-test (*\*\*\*P* < 0.0001). **c.** Comparison between the mutational spectra of the mutations identified in cSCC samples using WES and scRNA-seq data across the benchmarked algorithms. The cosine similarities between the mutational spectra derived from scRNA-seq and WES data are illustrated. **d.** Mutational rates for all samples using the somatic SNVs detected by scVar in cSCC and matched normal skin scRNA-seq data sets. **e.** Correlation between the mutation rate estimated using the mutations identified in WES and the mutations found in epithelial cells detected by scVar in the scRNA-seq data. The correlation was assessed using a linear regression model. Only genomic regions with sufficient sequencing depth available in both the WES and scRNA-seq datasets were included in this analysis. Mb, megabase.

### Validation of scVar using scRNA-seq and WES data

We further compared scVar with other methods using real data. Specifically, we employed matched WES and scRNA-seq data from eight cutaneous squamous cell carcinoma (cSCC) tumors[33]. Mutations characterized by high coverage in the WES data and concurrently detected in the corresponding scRNA-seq datasets were designated as the gold standard benchmark. This benchmark encompassed a total of 14,606 mutations located within regions of high sequencing coverage. These mutations, located in high-coverage areas, showed a positive correlation in VAF between WES and scRNA-seq (R = 0.52, *P* = 3.94×10^(-132), Supplementary Figs. 2a, 2b). Recall was operationally defined as the proportion of true mutations accurately identified by each computational tool, whereas precision was calculated based on mutations uniquely reported by the software but absent in both WES and scRNA-seq BAM files (see Methods). Benchmarking outcomes revealed that scVar significantly outperformed existing approaches. Compared to Monopogen, the second-best performing method, scVar achieved markedly higher recall rates (0.61–0.90 compared to 0.54–0.68; *P* = 0.00079, two-sided t-test) and F1 scores (0.75–0.92 versus 0.69–0.80; *P* = 0.0017), while consistently maintaining high precision values (0.89 to 0.98) across all evaluated conditions (Fig. 2b).

Subsequent performance evaluations were stratified by mutation subtypes and VAFs. Initially, mutations within the gold-standard dataset were categorized into six single-nucleotide substitution classes. scVar exhibited superior detection performance independent of subtype, achieving significantly higher recall rates (0.78–0.82) compared to the next-best performers, SComatic (0.44–0.48; *P* < 0.0004, two-sided t-test) and Monopogen (0.39–0.46; *P* < 0.0004), with no significant detection bias observed across single-nucleotide substitution subtypes (Supplementary Fig. 3a). Subsequently, we analyzed the VAF values of gold-standard mutations in WES tumor samples, comparing the recall rates of various software across different VAF intervals. In the critical ≤10% VAF range mutations, scVar achieved comparable performance to cellSNP. Furthermore, in the 10%–20% VAF range, scVar maintained superior recall value compared to SComatic, highlighting scVar’s efficacy in detecting low-frequency mutations (Supplementary Fig. 3b). Cosine similarity analysis of the mutation spectra further validated the correctness of mutation detection by scVar, demonstrating a high concordance with those identified by WES, with a cosine similarity score of 0.79 (Fig. 2c).

To validate scVar’s capability not only to detect a greater number of somatic mutations in cancer but also to resolve these mutations at the level of individual cell types, we analyzed mutation profiles from paired normal and tumor samples. This comparison revealed elevated mutation rates in tumor samples relative to matched normal tissues, with the exception of samples P2 and P10 (Fig. 2d). Additionally, mutation rates estimated from scRNA-seq data in epithelial cells exhibited a strong positive correlation with mutation burdens inferred from the WES data (R = 0.75, *P* = 0.032, linear regression; Fig. 2e), indicating that scVar facilitates precise quantification of mutation burdens at cell-type resolution.

Overall, these results demonstrate that scVar is a robust and effective tool for detecting somatic mutations in scRNA-seq data, outperforming established methods including cellSNP, Monopogen, and SComatic in terms of recall and F1 score. Its proficiency in accurately identifying mutations, particularly those with low VAF, alongside its consistent mutation spectrum concordant with WES data, underscores its reliability. Furthermore, scVar’s ability to quantify mutation burdens at the resolution of individual cell clusters offers novel insights into tumor heterogeneity, thereby advancing the understanding of cancer biology.

### Explore the heterogeneity at the sampling sites level and single-cell level in the non-small cell lung cancer (NSCLC) samples

#### Mutations Landscape of Different Sampling Sites

We analyzed five treatment-naïve, non-metastatic NSCLC patients undergoing curative-intent lobectomy[34]. Tumor specimens were dissected into core, mddle, and edge regions, alongside distal normal tissue located more than 5 cm from the tumor margin. These samples were dissociated into single-cell suspensions and subjected to droplet-based scRNA-seq. Due to insufficient yield (<500 cells) from the tumor core regions of patients P1 and P2, subsequent analyses were restricted to patients P3–P5: patient P3 (68-year-old male, stage IIIB lung adenocarcinoma), patient P4 (64-year-old female, stage IIB lung adenocarcinoma), and patient P5 (60-year-old male, stage IA3 large cell carcinoma). The scVar computational tool demonstrated superior performance in resolving spatial heterogeneity across these three NSCLC cases.

Clustering analysis of the mutation-cell binary matrix generated by scVar revealed distinct mutational landscapes among alveolar cells and T cells from patient P5 across different sampling sites (Figs. 3a, 3b). While transcriptional profiles exhibited limited spatial compartmentalization, mutational patterns displayed marked spatial segregation, indicating that genomic evolution occurs independently of transcriptional states. To thoroughly characterize spatial genomic heterogeneity, TMB was measured across distinct anatomical compartments within alveolar cells (tumor cells) and T cells. In both cell populations, the intermediate and peripheral regions demonstrated elevated TMB compared to the tumor core. Specifically, alveolar cells exhibited TMB values of 1.67 mutations per megabase (mutations/Mb) in the core, increasing to 2.33 and 2.37 mutations/Mb in the intermediate and edge regions, respectively. Similarly, T cells showed TMB values of 2.84 mutations/Mb in the core, rising to 3.99 and 3.80 mutations/Mb in the intermediate and edge compartments, respectively. These findings indicate the presence of distinct microenvironmental selection pressures across different tumor subregions (Fig. 3c).

**Figure 3.**
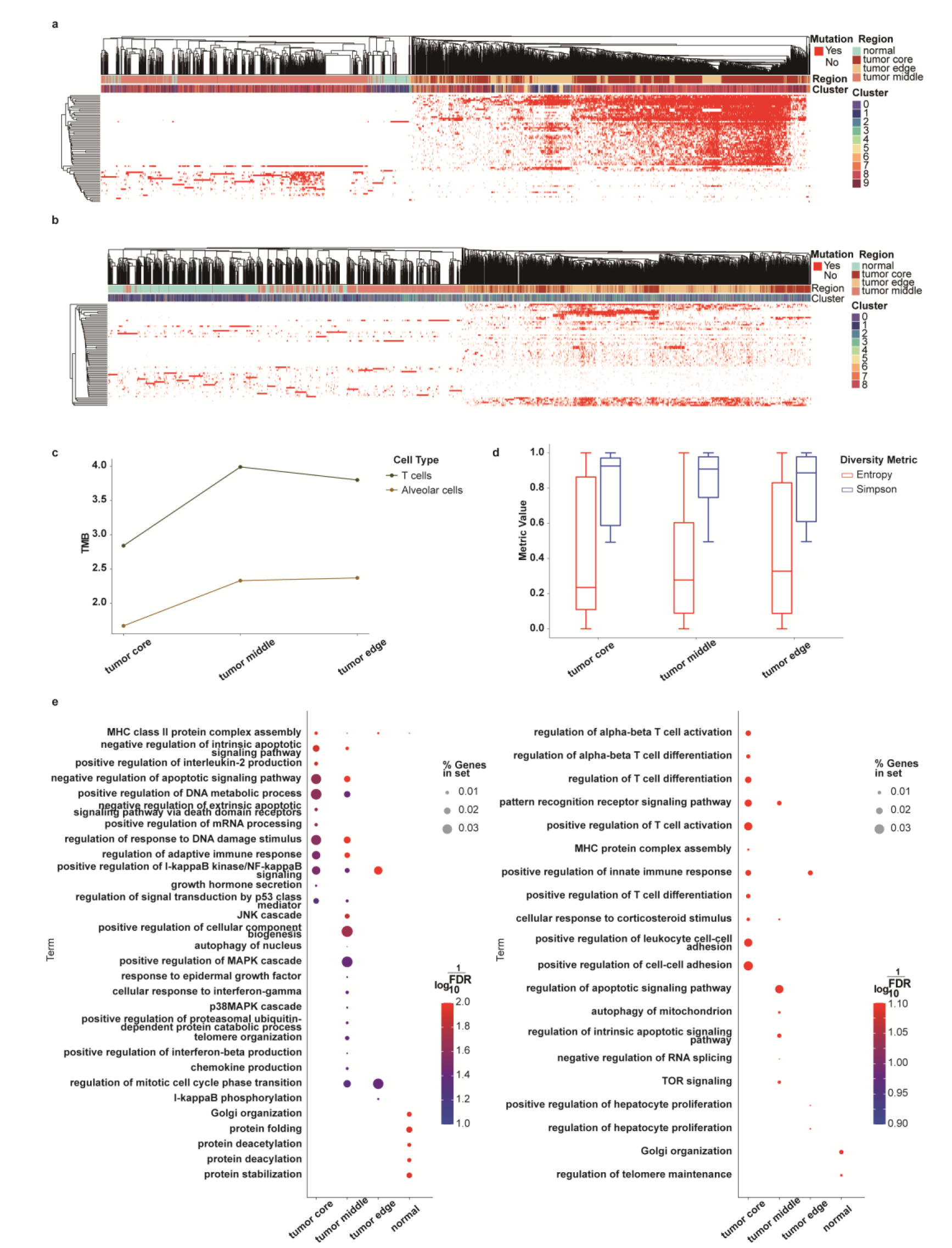
Analysis of spatial intra-tumor heterogeneity across tumor regions based on somatic mutations detected by scVar in scRNA-seq data. **a.** Hierarchical clustering of alveolar cells from all regions (represented in columns) based on somatic mutations (indicated in rows). Mutations identified in the cells are highlighted in red, while white signifies the absence of mutations. **b.** Hierarchical clustering of T cells across all regions (shown in columns) with somatic mutations (represented in rows). **c.** A line plot illustrating TMB in both alveolar cells and T cells across different sampling regions. **d.** Box plots displaying entropy and Simpson’s index for T cells at various tumor sites. **e.** GO enrichment results for mutated genes in alveolar cells (on the right) and T cells (on the left) across different sampling regions.

Quantitative assessment using entropy and Simpson’s diversity index revealed a progressive increase in T cell clonal diversity from the tumor core (mean entropy: 0.24; mean Simpson’s diversity index: 0.93) to the tumor edge (mean entropy: 0.33; mean Simpson’s diversity index: 0.89), with intermediate zones exhibiting transitional values (Fig. 3d). Comparable trends were observed in the other two patients (Supplementary Fig. 4). This gradient pattern aligns with microenvironmental pressure gradients, wherein hypoxic tumor cores favor dominant clones, while immune-enriched tumor margins support greater subclonal diversity, potentially mediated by differential exposure to microenvironmental factors and external selective pressures.

Functional partitioning was further elucidated through GO enrichment analysis of compartment-specific mutations. In alveolar cells, mutation-associated pathways exhibited distinct spatial heterogeneity across tumor subregions, with location-specific molecular signatures correlating with key oncogenic processes. Across the three tumor regions, pathways involved in the positive regulation of I*κ*B kinase (IKK)/ nuclear factor kappa B (NF-*κ*B) signaling were significantly enriched (adjusted *p*-values ranging from 0.005 to 0.037, hypergeometric test), consistent with prior findings that underscore NF-*κ*B’s established role in promoting tumor-associated inflammation and metastasis through sustained activation in lung carcinogenesis[35,36]. Both tumor core and intermediate regions exhibited enrichment of negative regulation of apoptotic signaling, suggesting systemic suppression of cell death mechanisms to sustain malignant survival. The unique enrichment of pathways associated with the positive regulation of mRNA processing in the tumor core may be attributable to its characteristic hypoxic and acidic microenvironment, as hypoxia has been shown to promote pre-mRNA splicing in the tumor microenvironment, leading to oncogene activation, tumor suppressor inactivation, metabolic adaptation, and angiogenic reprogramming[37]. In contrast, the tumor intermediate regions exhibited enrichment in multiple pathways linked to tumor microenvironment remodeling and metastasis, including: (1) epidermal growth factor response (EGF*R*), indicative of EGFR-mediated proliferative/migratory pathway hyperactivation characteristic of adenocarcinomas; (2) chemokine production, associated with epithelial-mesenchymal transition (EMT) induction and cancer stem cell niche formation[38]; and (3) co-enrichment of c-Jun-N-terminal kinase (JNK) cascade and p38 mitogen-activated protein kinases (p38MAPK) cascade, mechanistically linked to hypoxia-driven invasion via NOTCH4-mediated pathway crosstalk[39]. These activated pathways collectively enhance the invasive potential of lung adenocarcinoma cells. T cells isolated from the middle region of the tumor exhibited a specific enrichment in pathways associated with cancer development, particularly those regulating apoptotic signaling and intrinsic apoptotic pathways. Additionally, this region also demonstrated enhanced autophagy of mitochondria, which is essential for driving effector T cell differentiation through metabolic reprogramming[40]. This mechanism plays a significant role in the immunological regulation of tumor progression and the effectiveness of therapeutic responses[41] (Fig. 3e).

In summary, the application of scVar enabled detailed investigation of the genomic features in three NSCLC patients, thereby validating previously published observations at the single-cell level—an analytical resolution unattainable with conventional bulk genomic sequencing approaches.

### Mutations Landscape of Different Cell Types

The mutational landscape across distinct compartments of the tumor (core, middle, and edge) exhibited cell type-specific functional enrichment patterns that closely align with the intrinsic biological roles of these cells and their contributions to tumorigenesis. In patient P3, mutated genes within five predominant cell types—alveolar cells, B cells, fibroblasts, monocytes, and T cells—each comprising over 500 cells, were enriched in pathways related to the regulation of DNA damage response, mRNA splicing, and mRNA processing. These perturbations collectively contribute to genomic instability and aberrant transcriptional programs, which are hallmark features of cancer progression. Notably, alveolar cells demonstrated distinct enrichment of the EGFR signaling pathway and ErbB signaling pathway. The EGFR pathway, a well-established oncogenic driver in lung cancer, operates synergistically with other ErbB family members to activate critical intracellular signaling cascades that promote cell survival and proliferation[42]. Furthermore, alveolar cells showed functional enrichment in pathways governing mitochondrial gene expression, a process intimately linked to tumor progression. Dysregulated expression of mitochondrial respiratory genes— commonly suppressed in lung cancer tissues relative to normal tissues—has been directly correlated with tumor aggressiveness and poor prognosis. Notably, alveolar cells were also enriched for processes involved in surfactant homeostasis. Homeostasis of lung surfactants is closely related to alveolar cell function. Type II alveolar epithelial cells synthesize and secrete lipids and proteins of lung surfactants that maintain lung function[43]. While immune response-regulating signaling pathways were universally enriched among immune cells. Monocytes uniquely activated toll-like receptor 2 (TLR2) signaling and positively regulated NF-*κ*B-inducing kinase (NIK)/NF-*κ*B signaling, which are associated with pro-tumorigenic inflammation and immune evasion[44,45]. TLR2 activation via NF-κB has been reported to paradoxically promote proliferation in lung adenocarcinoma cells. Furthermore, monocytes displayed enrichment in pathways regulating phagocytosis and monocyte differentiation, which are closely related to monocyte function[46]. T cells are also enriched with antigen receptor-mediated signaling pathways and alpha-beta T cell activation associated with that cell type (Fig. 4a).

**Figure 4.**
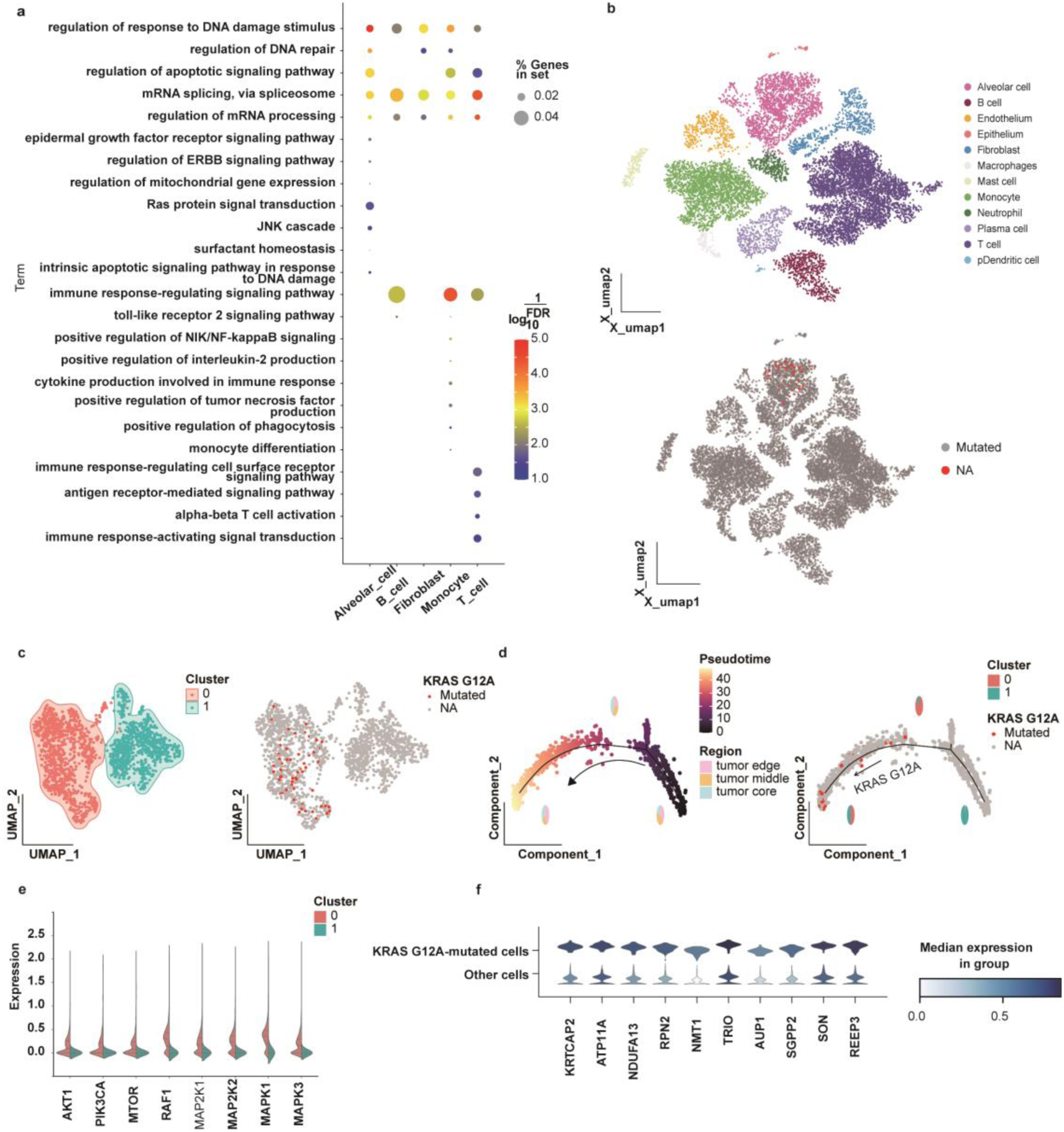
Analysis of intra-tumor heterogeneity across different cell types using somatic mutations detected by scVar in scRNA-seq data. **a.** GO enrichment analysis results of different cell types of mutant genes in three tumor samples from patient P3. **b.** The top inset presents the t-SNE plot of the 52,698 profiled cells annotated by associated cell type, while the bottom inset shows cells colored according to KRAS G12A mutation status as identified by scVar. **c.** UMAP visualization of alveolar cells. The left panel depicts the subclustering of alveolar cells, while the right panel shows the same cells colored according to KRAS G12A mutation status as identified by scVar. **d.** Pseudotime analysis of alveolar cells and proportion of cell sources at critical nodes shown in left. The right inset shows the position of *KRAS* G12A mutation cells in the trajectory and proportion of cell clusters at different nodes. **e.** Split violin plots illustrating the expression levels of genes associated with the RAF-MEK-ERK and PI3K-AKT-mTOR pathways in two alveolar cell subclusters. **f.** Stacked violin plots showing differentially expressed genes in *KRAS* G12A mutation cells compared to other alveolar cells.

Through analysis of mutations across distinct cell types, we observed that certain cancer driver mutations may exhibit cell type–specificity, thereby shaping their oncogenic potential within distinct cellular contexts. The *KRAS* gene, a pivotal member of the *RAS* oncogene family, represents the most frequently mutated oncogene across various malignancies[47]. Oncogenic *KRAS* mutations, including G12A, G12C, and G12D, result in constitutive activation of *KRAS*, driving aberrant signaling through pathways like MAPK and PI3K-AKT[48]. These mutations are associated with tumorigenesis, therapeutic resistance, and poor clinical outcomes in multiple cancers, including lung, colorectal, and pancreatic carcinomas[49,47,50]. The *KRAS* G12A mutation, although less common, is clinically significant and has been linked to reduced progression-free and overall survival in metastatic colorectal cancer, underscoring its role as a potent oncogenic variant[51]. According to the Cancer Hotspots database, 166 cancer patients, including six with lung cancer, harbor the *KRAS* G12A mutation[29,30]. In tumor samples from three NSCLC patients, 64 alveolar cells bearing *KRAS* G12A mutations were identified exclusively in patient P3 (Fig. 4b). Importantly, all *KRAS*-mutated cells were confined to the alveolar cell population, highlighting the cell type-specific nature of this oncogenic driver. Spatial analysis revealed an uneven distribution of *KRAS* G12A-mutated cells: 43 (67.2%) localized to the tumor core, 18 (28.1%) to the intermediate region, and 3 (4.7%) to the tumor periphery, suggesting enrichment within hypoxic or high-stress microenvironments.

Transcriptional profiling demonstrated that *KRAS* G12A-mutated alveolar cells possess distinct gene expression signatures relative to their wild-type counterparts, indicative of mutation-driven transcriptional reprogramming. All *KRAS* G12A-mutated cells clustered uniquely within Cluster 0, exhibiting clear separation from wild-type alveolar cells in UMAP visualization (Fig. 4c). Pseudotime trajectory analysis positioned *KRAS* G12A-mutated alveolar cells predominantly at the terminal branch of the developmental continuum (Fig. 4d), suggesting a later differentiation stage compared to wild-type cells. This observation is similar to previous findings in KRAS G12D-mutated alveolar cells[52].

Canonical *KRAS* signaling activates the RAF-MEK-ERK cascade and also modulates the PI3K-AKT-mTOR pathway, a critical axis regulating cell proliferation, differentiation, apoptosis, and glucose metabolism, with profound implications for tumorigenesis and therapeutic resistance. Comparative analysis revealed that clusters harboring the *KRA*S G12A mutation exhibited elevated expression of genes associated with both the RAF-MEK-ERK and PI3K-AKT-mTOR pathways relative to control clusters (Fig. 4e), suggesting hyperactivation of these oncogenic signaling networks[47]. Differential gene expression analysis identified multiple genes implicated in lung cancer and other malignancies as significantly upregulated in *KRAS* G12A-mutant cells compared to other alveolar cells (Fig. 4f). Notably, *ATP1A1*, *NMT1*, *IFITM1*, and *BPTF* were markedly overexpressed. These genes have established associations with lung cancer pathogenesis. Specifically, *BPTF* promotes tumor growth and correlates with poor prognosis in lung adenocarcinoma[53], while *IFITM1* drives tumor cell metastasis—effects demonstrably inhibited by its silencing in lung cancer models[54]. Additionally, KRTCAP2 displays the most pronounced differential expression, with established roles in hepatocellular carcinoma (HCC)[55]. Other cancer-relevant differentially expressed genes (DEGs) include the recognized oncogene *RPN2*, which enhances the malignant properties of tumor cells and affects multidrug resistance[56]; *REEP3*, overexpressed in pancreatic cancer and correlated with reduced overall survival and disease-free survival[57]; The ARF family is generally highly expressed across pan-cancer, and *ARF5* is significantly associated with poor prognosis in HCC patients[58]. This comprehensive profile underscores the multifaceted oncogenic signatures associated with *KRAS* G12A mutations.

In summary, relative to conventional single-cell transcriptome analyses, scVar enables the integration of somatic mutation data with transcriptomic profiles at single-cell resolution. This approach facilitates a more comprehensive investigation of the heterogeneity among diverse cell types and subpopulations within tumors. Consequently, it yields novel insights into tumor heterogeneity and presents promising opportunities for the development of targeted therapeutic interventions.

## Discussion

This study presents scVar, a comprehensive framework capable of detecting SNVs from 10X scRNA-seq data and performing joint analyses of gene expression and genetic variation at single-cell resolution. Building on softwares established for bulk transcriptomics, scVar extends mutation detection to 10x Genomics scRNA-seq data. Importantly, scVar demonstrates enhanced sensitivity in identifying somatic mutations, particularly low-frequency variants, without the necessity for paired normal samples or predefined cell-type annotations. The software further incorporates comprehensive downstream analytical modules that facilitate the exploration of heterogeneity among distinct cellular subpopulations. These modules include the characterization of mutational signatures, quantification of tumor mutational burden and clonal diversity, and functional enrichment analyses of mutated genes. Through the joint analysis of mutational and transcriptomic features, scVar can reveal differential gene expression programs in mutated cells, delineate their trajectories within clonal and developmental lineages, and correlate transcriptional heterogeneity with genetic variation at single-cell resolution, thereby providing a new perspective for the study of tumor heterogeneity. SNVs, particularly low-frequency variants, often play an essential role in tumor evolution and heterogeneity. In benchmarking with both simulated datasets and eight cSCC tumors with matched WES and scRNA-seq, scVar consistently outperformed comparator methods, achieving higher recall and F1 scores with enhanced sensitivity to rare mutations.

Integrating genomic and transcriptomic information at single-cell resolution facilitates direct mapping of genotype-to-phenotype relationships, thereby advancing the understanding of tumor heterogeneity. scVar can be seamlessly incorporated into standard 10X Genomics workflows, enabling the concurrent analysis of gene expression and mutation data, which enhances mutation-level insights and broadens its practical applicability. Application of scVar to real datasets has enabled detailed investigations of mutational characteristics across diverse cell populations, facilitated the examination of transcriptional programs in mutated cells such as those harboring oncogenic KRAS mutations, and supported the tracking of mutated cells along cellular trajectories, thereby providing deeper insights into intratumoral heterogeneity.

Despite these advances, several limitations warrant consideration and motivate future developments. First, somatic mutations present in a limited subset of single cells may evade detection by WES or WGS, resulting in reduced cosine similarity between the mutation spectra derived from scVar and WES data. Second, current scRNA-seq technologies face challenges including data sparsity, elevated dropout rates, and inadequate gene expression levels. The lack of full-length transcript coverage in platforms such as 10X Genomics platform further complicates data completeness and mutation detection accuracy. To overcome these issues, future developments of scVar will focus on enhancing compatibility with full-length single-cell transcriptome protocols and long-read sequencing technologies, thereby improving variant identification and enabling the detection of alternative splicing events. Additionally, systematic sequencing artifacts and platform-specific errors continue to contribute substantially to false-positive variant calls. Consequently, the development of a Panel of Normals (PON) tailored for 10X scRNA-seq data, which catalogs recurrent artifact sites and models background noise, represents a promising approach to enhance variant calling accuracy. Moreover, copy number variations (CNVs) and structural variations (SVs) are critical contributors to the genomic instability characteristic of cancer cells, influencing tumorigenesis, tumor behavior, and therapeutic response. To address these challenges, future iterations of scVar aim to incorporate CNV and SV detection capabilities, thereby enabling a more comprehensive mutational analysis and enhancing the overall applicability and analytical robustness.

## Conclusions

scVar is an integrated workflow designed to detect SNVs from 10x Genomics scRNA-seq data and to perform joint analyses of gene expression and genetic variation at single-cell resolution. Evaluation using simulated datasets as well as matched WES and scRNA-seq data demonstrates that scVar outperforms existing methods in identifying somatic mutations from scRNA-seq, exhibiting notably enhanced sensitivity for variants with low VAF. Application of scVar to cancer datasets enables the seamless integration of somatic mutation detection with transcriptomic profiles at the single-cell level, thereby providing more nuanced insights into tumor heterogeneity.

## Methods

### Detection of somatic mutations in single-cell data sets using scVar Initial read mapping and scRNA-seq expression analysis

During the initial stage of the scVar protocol, sequenced reads are aligned to the reference genome using the count tool of Cell Ranger[59] Count (v.7.1.0) under default parameters. To reduce the impact of cell-free RNAs that constitute background noise, scVar employs soupX[60] (R v.4.1.2 and soupX v.1.6.2) to estimate and remove ambient RNA contaminations.

A seurat object is created using CreateSeuratObject in Seurat[61] (v.4.3.0.1) based on the cleansed output from soupX with the parameters min.cells and min.features set to 3 and 10 respectively. Cells that contain greater than 10 and less than 15,000 expressed genes, fewer than 50% ribosomal transcripts, and fewer than 10% mitochondrial transcripts are saved. The data matrix undergoes a standardization process through NormalizeData, utilizing a scale.factor of 1e4. Top 2,000 variable features are identifird by FindVariableFeatures using variance stabilization transformation (“vst”). After further scaling data by ScaleData, Principal component analysis (PCA) is performed with the variable features by RunPCA. Unsupervised graph-based clustering is presented by FindNeighbors with 30 reduction dimensions and FindClusters with the resolution set to 0.7. SingleR[62] (v.1.8.1) is used to detect cell types based on Human Primary Cell Atlas data as the reference.

### Mutation detection and annotation

To minimize the impact of low-quality cells on subsequent analyses, the reads are extracted from BAM files to create new BAM files based on the barcodes of high-quality cells identified after quality control. First, scVar separates the BAM files by chromosome using the split tool from Bamtools[63] (v.2.5.1), and employs the pysam package (Python v.3.11.4) to extract the reads corresponding to high-quality cells identified by the ’CB’ tag. The Multiprocessing package is utilized for parallel processing.

The extracted BAM files are sorted and duplicates are marked using the SortSam, BuildBamIndex, and MarkDuplicates functions from Picard (https://broadinstitute.github.io/picard/) (Java v.1.8.0_382 and Picard v.2.25.5). The SplitNCigarReads tool from GATK[24–26] (v.4.2.2.0) is used to split reads containing Ns. To mitigate systematic bias, scVar utilizes GATK BaseRecalibrator and ApplyBQSR to recalibrate base quality, referencing the high-confidence SNPs from the 1000 Genomes Project (1000G) Phase 1

(1000G_phase1.snps.high_confidence.hg38.vcf.gz), as well as Mills and 1000G gold standard indels (Mills_and_1000G_gold_standard.indels.hg38.vcf.gz), and dbSNP version 138 (downloaded as part of the GATK bundle). Candidate mutations are identified using GATK HaplotypeCaller. Due to the relatively low data coverage associated with 10X, scVar specifically sets the ploidy parameter to 40 to facilitate the detection of low-frequency mutations. Both the VEP[27] (v.107) and Annovar[28] (Perl v.5.34.0) are used to prioritize and annotate the genomic variants and assess their effects. Concurrently, scVar compiles several clinical databases using Vcfanno[64] (v.0.3.3), including DOCM[65], CancerHotspot[29,30], CIVIC (https://civicdb.org/welcome), COSMIC[31], CGI[66,67], and ClinVar[32], and incorporates these annotations into the INFO field of the VCF file.

### Mutation genotype assignment and filter

To investigate mutations at the single-cell level, scVar quantifies reference and alternate reads for individual cells at each mutation site leveraging the pysam package (v.0.23.0). This process utilizes the unique molecular identifier (UMI) and ’CB’ tags embedded in sequencing reads. To ensure robust genotyping, users can configure critical quality thresholds: (1) a minimum read count per barcode to exclude cells with insufficient sequencing coverage, (2) a minimum mapping quality threshold to filter ambiguously aligned reads, and (3) a minimum base quality score to enhance confidence in base calling. To optimize computational efficiency, the mutation file is partitioned by chromosome, with each chromosomal dataset further subdivided into multiple .tbl files based on user-defined thread specifications. These segmented files are processed in parallel using the multiprocessing package, enabling scalable analysis across large mutation datasets. Ultimately, scVar presents the results detailing the counts of reference and alternate reads for all cells at different mutation sites. A mutation is considered present in a cell if at least one alternate read supports it. To reduce noise, scVar retains only mutations that are present in at least two cells. Mutations catalogued in gnomAD[68] (v.4.1) or ExAC[69] (v.0.3) with a population allele frequency greater than 1%, as well as those identified within a PON constructed from WGS data of the 1000 Genomes Project (https://gatk.broadinstitute.org/hc/en-us/articles/360035890631-Panel-of-Normals-PON-), are systematically excluded from analysis. Additionally, users can set a maximum VAF threshold to refine mutations further, excluding variants exceeding a user-defined VAF limit to reduce potential germline contamination.

Finally, scVar generates a sparse cell-by-mutation matrix where -1 indicates no coverage, 0 denotes wild-type, and 1 represents mutation. The chunk processing is employed to optimize memory efficiency. For multi-omics integration, results are structured as a MuData object containing transcriptomic data in the rna modality and mutation data with genomic annotations in the mutation modality, enabling comprehensive joint analysis of genomic variants and gene expression profiles.

### Downstream analysis in scVar

#### Mutation signature, TMB, Entroy and Simpson’s index analysis

scVar consolidates mutations within each cell type to comprehensively analyze the mutation processes across diverse cellular populations. Initially, scVar computes the proportion of mutations occurring in each of the 96 trinucleotide contexts and evaluates the similarity of mutational signature derived from COSMIC[31] (v.3.3.1) for both the overall results and cell type-specific subsets, employing deconstructSigs[70] (v.1.8.0). To assess tumor burden, scVar incorporates mosdepth[71] to assess the length of the genomic regions covered by the BAM file, focusing specifically on the autosomes and the X and Y chromosomes. Subsequently, based on the mutation results, scVar quantifies the number of non-synonymous mutations from the mutation results, including Frame_Shift, Splice_Site, Translation_Start_Site, Nonsense_Mutation, Nonstop_Mutation, and Missense_Mutation types. The tumor burden for each patient is subsequently calculated using the following formula.

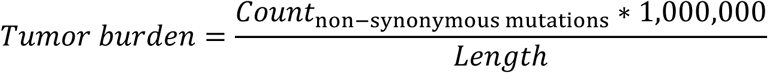

To quantify intertumoral genomic heterogeneity, scVar calculates Shannon entropy and Simpson’s diversity index for each mutation. Entropy(*H*) is calculated according to the following formula:

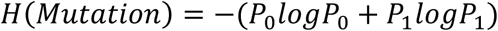

where *P*_0_ denotes the probability of no mutation and *P*_1_ denotes the probability of mutation.

Simpson’s diversity index(*D*) is calculated according to the following formula:

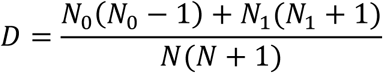

where *N*_0_ equals the number of non-mutated cells, *N*_1_ equals the number of mutated cells and *N* corresponded to the total number of cells, including both mutated and non-mutated populations.

### Cell type-specific mutations detection

To examine the relationship between genetic mutations and cellular functional heterogeneity, scVar detects mutations that are specific to distinct cell types. For each identified mutation, the counts of mutated and non-mutated cells are determined across predefined cell types or subclusters. A two-sided Fisher’s exact test is then performed using the fisher_exact function from the Python package scipy.stats (version 1.15.2) to assess the enrichment of mutations within specific cell populations. Subsequently, *p*-values are adjusted for multiple comparisons using the Benjamini-Hochberg (BH) method implemented through the multipletests function from the statsmodels package (version 0.14.4). A comparable statistical approach is employed at the gene level: for each mutated gene, counts of mutated and non-mutated cells are stratified by cell type or subcluster, followed by the application of Fisher’s exact test and subsequent multiple testing correction. This approach enables the systematic identification of mutations and genes exhibiting significant specificity to particular cell types, thereby establishing a connection between genetic variation and functional cellular heterogeneity.

### Cluster cells based on mutations

The final output of scVar consists of a binary matrix delineating cells by mutations, which forms the foundation for mutation-driven cell clustering. Two primary feature extraction methodologies are utilized: (1) direct calculation of variance on the binary matrix to identify mutations with high variability across cells, and (2) variance-based feature selection following TF-IDF normalization, intended to mitigate noise arising from the matrix’s pronounced sparsity. The TF-IDF transformation is performed using the TfidfTransformer function from the scikit-learn library (version 1.5.2).

Subsequent to feature selection, pairwise Jaccard distances between cells are computed to quantify dissimilarities in their mutational profiles. The hierarchical clustering framework implemented in scVar supports multiple linkage methods, including the unweighted pair-group method with arithmetic mean (UPGMA) and the weighted pair-group method with arithmetic mean (WPGMA), to construct dendrograms. Finally, clustering results are visualized via the pheatmap function from the ComplexHeatmap package (version 4.4.4), which incorporates dendrograms to highlight cell subpopulations distinguished by unique mutational signatures.

### Oncogenic pathway and GO function enrichment analysis

scVar performs comprehensive enrichment analysis for mutated genes both across the entire cell population and within distinct cell types. The analysis focuses on functionally significant mutation types including: Frame_Shift, Splice_Site, Translation_Start_Site, Nonsense_Mutation, Nonstop_Mutation, Missense_Mutation, as well as variants located in intergenic regions (IGR), 5’ untranslated regions (5’UTR), and 3’ untranslated regions (3’UTR). Mutations classified as non-synonymous but lacking definitive functional implications are excluded from the analysis. For the investigation of oncogenic pathways, scVar employs the OncogenicPathways function from the maftools package (version 2.12.0) in R, with the resulting data visualized through heatmaps generated using the seaborn library (version 0.13.2) in Python. GO enrichment analysis is performed utilizing the enrichGO function from the clusterProfiler package (version 4.4.4), referencing the org.Hs.eg.db database. The analysis permits user specification of statistical significance thresholds via adjustable pvalueCutoff and qvalueCutoff parameters, with *p*-values corrected for multiple comparisons using the BH method. Visualization of enrichment outcomes is achieved through point plots created with matplotlib (version 3.10.1).

### Pseudotime analysis

To investigate the distribution of mutated cells along differentiation trajectories, scVar implements time-series analysis using Monocle[72] (v2.24.0), with default parameters. Genes are included in the analysis only if they are expressed in a minimum of 10 cells. The setOrderingFilter function is applied to sort cells based on differentially expressed genes (qval < 0.01) among cell subsets. The reduceDimension function is used to generate traces, employing the DDRTree method. Finally, the trace plot is produced using ggplot.

The core workflow of scVar is managed through the Snakemake pipeline management system (version 3.13.3), enabling automated and reproducible execution of analytical processes. Subsequent analysis components are developed as standalone scripts, providing flexibility for tailored customization of individual analytical tasks. scVar generates an extensive, interactive HTML report dynamically rendered using Vue.js, which incorporates visualizations, statistical summaries, and relevant metadata. The software is fully containerized via Docker technology, ensuring a platform-independent environment that supports computational reproducibility across various computing systems.

### Method of simulating data

To rigorously assess the performance of variant calling software, we implemented a benchmark using simulated data. Initially, mutation site selection was performed as follows: for each input BAM file, genome-wide coverage profiles were generated using mosdepth[71]. Candidate insertion sites (n=5000 per sample) were then identified from genomic regions exhibiting sequencing depths of at least 10X, utilizing a multi-step filtering strategy implemented via pysam. This filtering procedure involved: (a) exclusion of population variants catalogued in gnomAD[68] (v.4.1) or ExAC[69] (v.0.3) with a population frequency exceeding 1%, as well as variants present in a PON generated from WGS data of the 1000 Genomes Project, (b) retention of sites with canonical reference nucleotides (A/T/C/G), and (c) enforcement of stringent quality criteria, specifically mapping quality ≥200 and base quality ≥35. The distribution of mutations across chromosomes was proportionally allocated based on chromosome length, and multiprocessing techniques were employed to optimize computational efficiency. Subsequently, BAM file modification was conducted by introducing controlled mutagenesis at the selected candidate sites. Initially, all reads were reverted to their reference-aligned states using pysam. Site-specific variants were then incorporated such that the allele fractions of the modified reads precisely matched predetermined VAFs of 10%, 20%, 30%, 40%, and 50%, resulting in five simulated samples per input file. The performance of four variant callers—cellSNP, Monopogen, SComatic, and scVar—was assessed using the F1-score metric. Precision and recall were systematically computed by comparing the ground-truth mutations with the variants identified by each caller across all VAF levels.

### Valuation with WES

To validate the performance of the scVar software, we leveraged the paired WES and scRNA-seq datasets, employing mutations identified from the WES data as the reference gold standard. The WES data were aligned to the human reference genome using BWA MEM[73] (v.0.7.17). Subsequent processing of the resulting BAM files included duplicate removal and base quality score recalibration in accordance with the GATK Best Practices workflow. SNVs from WES data were identified using Strelka2[74] with default parameters, incorporating matched normal samples as germline controls.

Genomic regions were then filtered to retain those with a minimum sequencing depth of 1X in the WES data and at least 10X in the scRNA-seq data, as determined by mosdepth[71]. To define a set of gold standard mutation candidates, overlapping regions between these datasets were determined via the Intersect function of Bedtools[75]. Regenotyping was performed on candidate mutations present in both the WES and scRNA-seq BAM files. Gold standard mutations were defined based on the following criteria: absence of alternative allele reads in the WES normal sample, presence of at least one alternative allele reads in the WES tumor sample, and more than two alternative allele reads in the scRNA-seq tumor sample.

For performance evaluation, true positives (TP) were defined as mutations detected by scVar that were included in the established true mutation set. False negatives (FN) comprised mutations within the true set that scVar failed to identify. False positives (FP) were mutations uniquely reported by scVar but subsequently determined to be absent upon regenotyping in both WES and scRNA-seq BAM files.

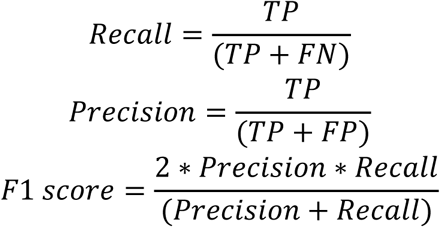

### Methods of analyzing real data

#### Single-cell gene expression quantification and determination of the major cell types

Raw gene expression matrices were generated for each sample using Cell Ranger Count and SoupX, and merged using Scanpy[76] (v.1.10.2) concat method. Cells were filtered to retain those expressing more than 10 and fewer than 15,000 genes, with ribosomal transcript content below 50% and mitochondrial transcript content below 10%. Normalization was conducted by scaling the total counts per cell to 10,000 transcripts (normalize_total(target_sum=1e4)) followed by log(1+x) transformation. Highly variable genes (HVGs) were identified using Seurat’s variance-stabilizing method (highly_variable_genes(n_top_genes=2000, flavor=’seurat’)), selecting the top 2,000 genes for downstream analyses. Prior to PCA, data were scaled with a maximum value of 10 (scale(max_value=10)). PCA was conducted with default parameters, retaining the first 50 principal components. A k-nearest neighbor graph was constructed using 10 neighbors and 50 principal components (neighbors(n_neighbors=10, n_pcs=50)), followed by UMAP embedding with a minimum distance parameter of 1 (umap(min_dist=1)). Cell clustering was performed using the Leiden algorithm at a resolution of 1, implemented via Leidenalg (leiden(resolution=1, random_state=0, flavor="leidenalg", key_added="leiden_res1")). UMAP visualizations were generated using the utils.embedding function from the Omicverse[77] package (v.1.6.7). Cell type annotation was conducted by integrating marker genes from the CellMarker database[78] (CellMarker Augmented (2021)) and markers reported in the literature. For annotation based on database-derived marker genes, pathway enrichment analysis was carried out using the ov.single.pathway_aucell_enrichment function from the Omicverse package, which mapped gene sets from a predefined pathway dictionary (pathways_dict) to cellular activities utilizing eight parallel workers (num_workers=8). Differential gene expression analysis was subsequently executed using sc.tl.rank_genes_groups with the Wilcoxon rank-sum test (method="wilcoxon"), grouping cells according to Leiden clustering results. The top three differentially expressed genes per cluster were visualized as a dot plot through sc.pl.rank_genes_groups_dotplot, generated by variance-scaled expression values (standard_scale=’var’).

### Subclustering and trajectories analysis of alveolar cells

To delineate subclusters within alveolar cells, we conducted a focused reanalysis of cells classified as alveolar. Initially, raw count data specific to alveolar cells were extracted. Subsequent data preprocessing involved normalization and logarithmic transformation. HVGs were identified employing the Seurat variance-stabilizing approach (sc.pp.highly_variable_genes) with the following parameters: min_mean = 0.0125, max_mean = 3, min_disp = 0.5, and flavor set to ’seurat’. PCA was then performed using default settings. A k-nearest neighbor graph was constructed utilizing 10 neighbors and 40 principal components (sc.pp.neighbors(n_neighbors=10, n_pcs=40)). UMAP embedding was subsequently applied with a minimum distance parameter of 0.5 (sc.tl.umap(min_dist=0.5)). To identify transcriptionally distinct subpopulations, Leiden clustering was implemented at a resolution of 0.1 (sc.tl.leiden). Finally, pseudotime analysis was conducted across all alveolar cells to infer dynamic cellular trajectories. To investigate pathway activity within alveolar subclusters, we compared the expression patterns of genes associated with the RAF– MEK – ERK cascade and the PI3K – AKT – mTOR signaling pathway. Visualization of gene expression level was performed using violin plots generated with the seaborn (sns.violinplot) package. Differential expression analysis was conducted between cells harboring the *KRAS* G12A mutation and those without detectable *KRAS* G12A mutation, using the sc.tl.rank_genes_groups function in Scanpy with the t-test method. From the top 20 genes ranked in this analysis, we concentrated on those implicated in tumorigenesis and cancer progression, particularly oncogenes and driver genes previously documented in lung cancer and other malignancies. Notable genes identified included KRTCAP2, ATP1A1, RPN2, NMT1, REEP3, IFITM1, ARF5 and BPTF.

## Availability of data and materials

scVar is available on GitHub https://github.com/WeiZhaoLuck/scVar and Biocode https://ngdc.cncb.ac.cn/biocode/tool/BT008000 <u>at National Genomics Data Center</u>[79]. The code is released under the version 3 of the GNU General Public License. Matched WES and scRNA-seq datasets from eight cutaneous squamous cell carcinoma (cSCC) tumors were retrieved from the NCBI Gene Expression Omnibus database under accession numbers GSE144237 (https://www.ncbi.nlm.nih.gov/geo/query/acc.cgi?acc=GSE144237) and GSE144240 (https://www.ncbi.nlm.nih.gov/geo/query/acc.cgi?acc=GSE144240). Public single-cell lung cancer data were downloaded from ArrayExpress under accession E-MTAB-6149 (https://www.ebi.ac.uk/biostudies/arrayexpress/studies/E-MTAB-6149/files).

## Acknowledgements

We would like to thank Yingke Ma, Song Wu, Tong Jin, Mochen Zhang, and anonymous reviewers for helpful feedback and discussions.

## Contributions

Y.B., X.Z., and W.Z. designed the project. W.Z. and X.Z built the software. W.Z. conducted the software benchmark, application, and analysis. Y.B., X.Z., and W.Z. wrote the manuscript. All authors contributed to editing the manuscript. All authors read and approved the final manuscript.

## Ethics declarations

### Ethics approval and consent to participate

Not applicable.

### Consent for publication

Not applicable.

## Additional information

### Supplementary Information

#### Supplementary Note

To rigorously assess the efficacy of mutation detection tools in identifying low-frequency variants from 10X scRNA-seq data, we developed a simulation framework grounded in real scRNA-seq datasets. This framework involved the computational generation of hybrid tumor samples by mixing sequencing reads from two distinct individuals (referred to as sample1 and sample2) at predetermined ratios (10%:90%, 20%:80%, 30%:70%, 40%:60%, and 50%:50%).

The simulation process comprised four principal stages. Initially, genomic regions exhibiting overlapping coverage between the two samples were identified using Bedtools (employing the bamtobed, merge, and intersect functions). Subsequently, reads corresponding to these shared regions were extracted from each BAM file utilizing Samtools. To simulate varying mutation frequencies, differential downsampling of the original BAM files was performed via Picard’s DownsampleSam function, retaining 10–50% of reads for sample1 and 90–50% for sample2, followed by merging with Samtools.

For the establishment of a gold-standard mutation set, a stringent filtering pipeline was applied: (1) initial variant calling was conducted using Strelka2 (version 2.9.10), requiring variants to exhibit 100% VAF and coverage exceeding five reads in both samples; (2) homozygous germline mutations specific to sample1 were selected as candidate variants; (3) regenotyping validation across all samples was performed to retain only mutations demonstrating complete allele segregation (i.e., 100% alternative alleles in sample1 and 100% reference alleles in sample2); and (4) final VAF thresholding was applied, allowing a ±2% deviation from the expected downsampling ratio to define the benchmark mutation set.

We conducted a systematic comparison of six mutation detection methodologies, including the GATK Best Practices workflow (incorporating Picard MarkDuplicates, SplitNCigarReads, BaseRecalibrator, ApplyBQSR, and HaplotypeCaller with ploidy set to 40), Strelka2 (v2.9.10), and VarScan (v2.3.9). Recall rates were computed as the ratio of TP to the sum of true positives and false negatives (TP + FN), where true positives corresponded to gold-standard variants detected by each tool, and false negatives represented benchmark mutations that were not identified.

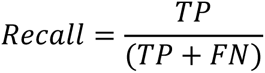

Notably, the optimized GATK pipeline exhibited superior performance across all mixing ratios, achieving recall rates ranging from 84.5% to 96.3%, in contrast to 62.1% to 89.4% observed for alternative tools (Supplementary Figure 1a). This performance advantage persisted when mutations were stratified by substitution type (C>A, C>G, C>T, T>A, T>C, T>G), with GATK and Strelka2 demonstrating comparable detection rates across all categories (Supplementary Figure 1b). The demonstrated robustness of GATK, particularly at low mutation frequencies (10–30% mixtures), informed its selection as the primary variant caller within the scVar analytical framework.

