## Supplementary figures and images for "Joint analysis of expression and variation at single cell resolution by scVar"

### Supplementary Fig. 1

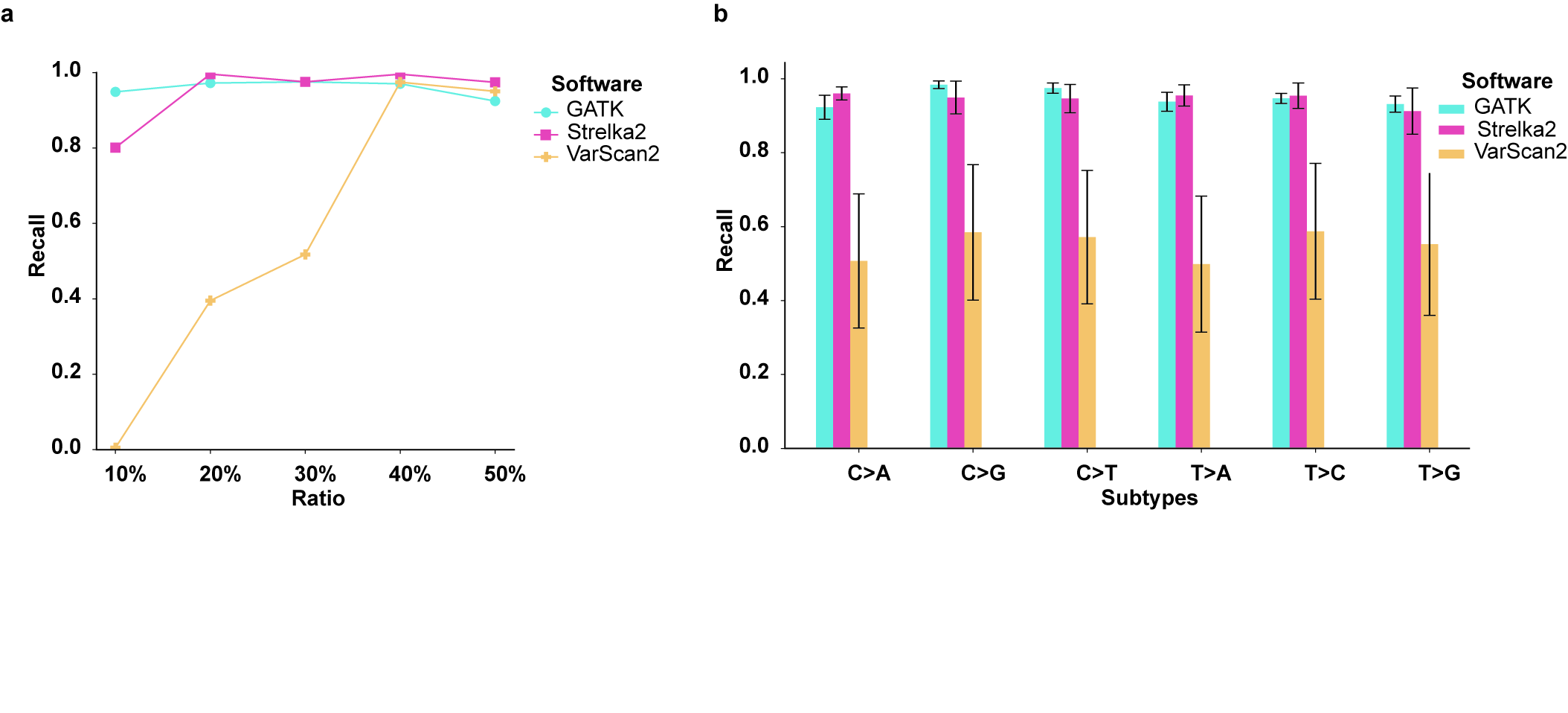

### Supplementary Fig. 2

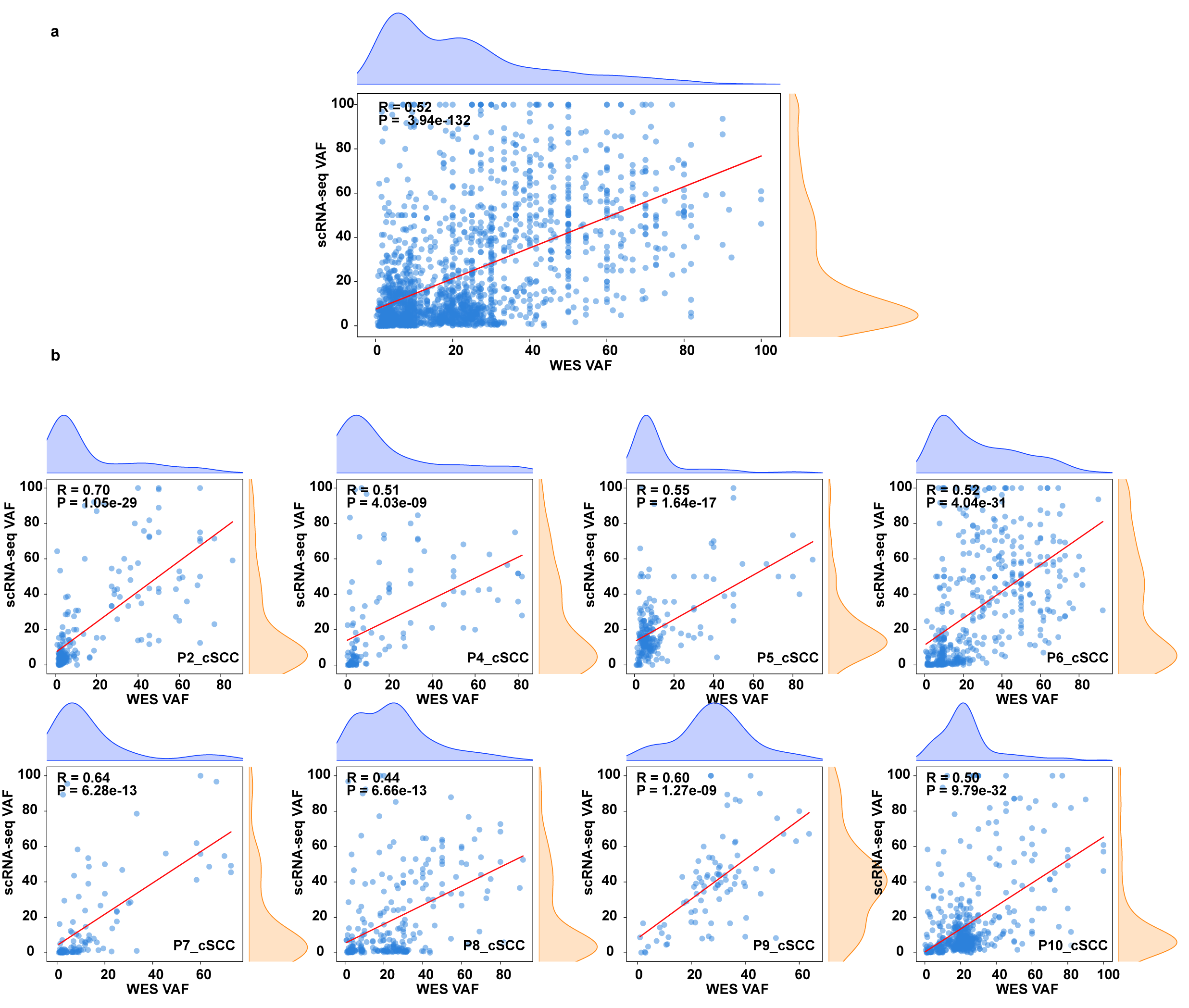

### Supplementary Fig. 3

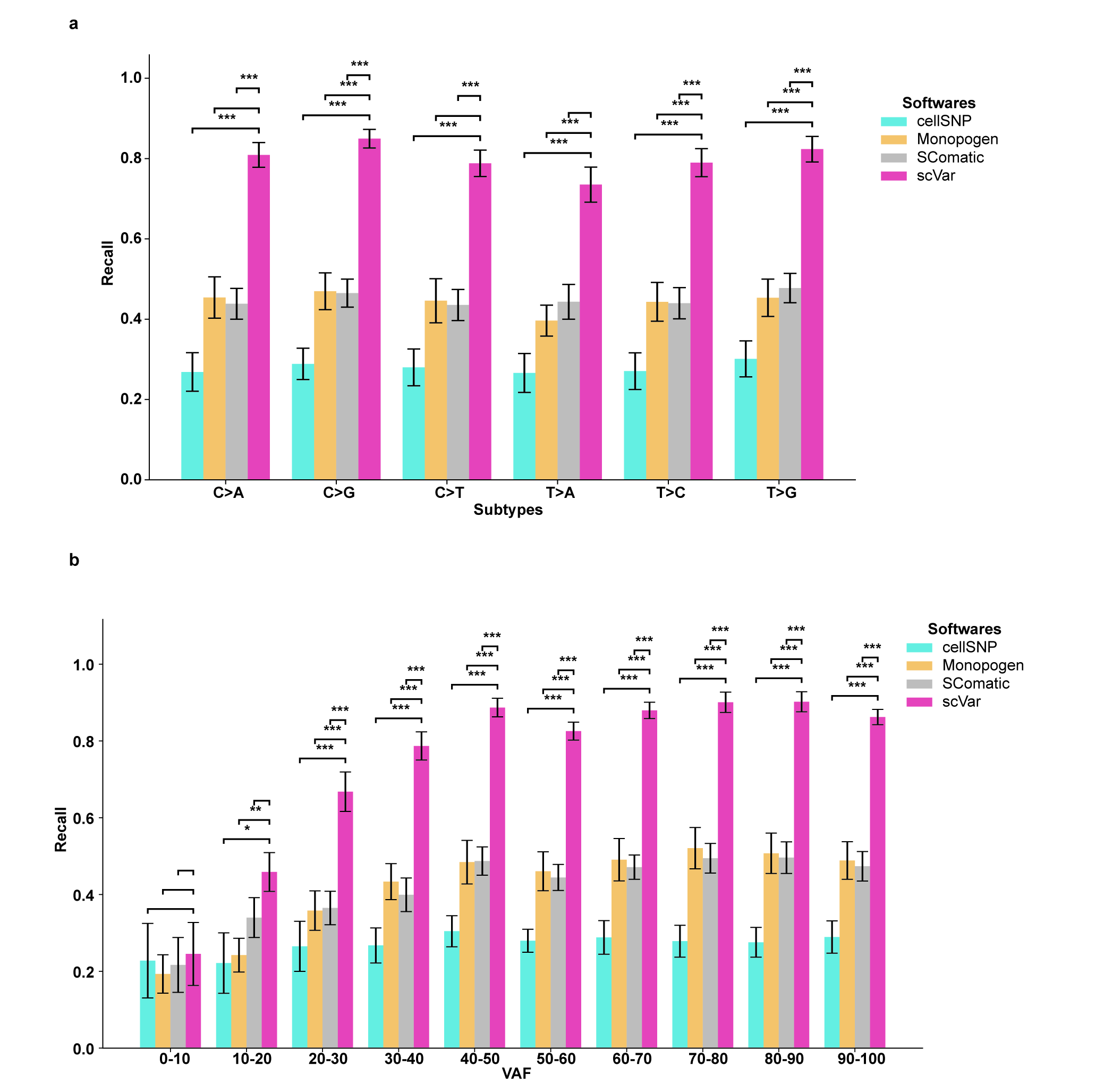

### Supplementary Fig. 4

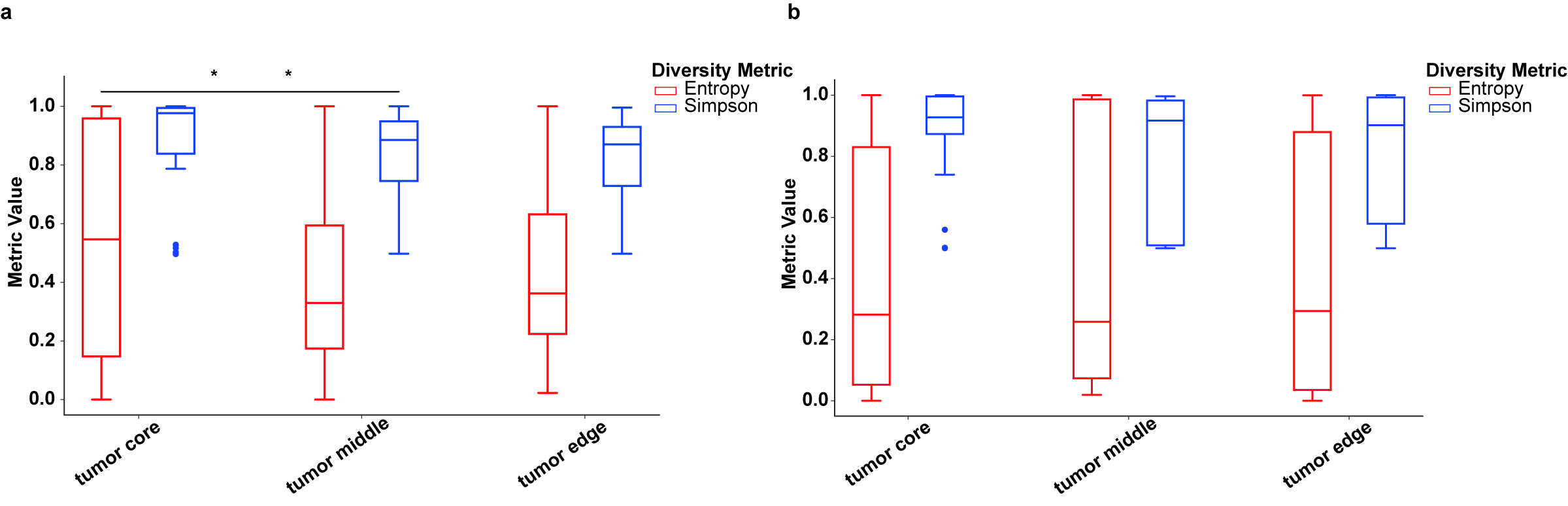
